# Comparison between Canine and Porcine Models of Chronic Deep Venous Thrombosis

**DOI:** 10.1101/2023.02.20.529322

**Authors:** Chuang Wang, Tao Tang, Sheng-Lin Ye, Xu-Dong Jiang, Guang-Yuan Xiang, Lun Xiao, Lu-Lu Wang, Tian-Ze Xu, Bin Song, Nan Hu, Xiao-Long Du, Xiao-Qiang Li

## Abstract

Venous thromboembolism(VTE) refers to deep venous thrombosis(DVT) and pulmonary embolism(PE), which is a worldwide problem and has very high morbidity and mortality. The research of venous thrombosis involves pathogenesis, pathophysiology, and therapeutic attempt. Alternative experimental animal models have changed dramatically over the past 20 years, from small murine models to large ones. Larger animal models offer more options and are more consistent with human physiology. This article aims to induce chronic deep venous thrombosis in the left iliac vein in canine and porcine models and compare these two models. We think that feasible large animal models can better translate the results of therapeutic research into clinical application.

## Introduction

The use of animal models to construct venous thrombosis is an essential means to study the occurrence, development, and treatment of thrombosis. Most of the models are based on small rodents to construct acute or subacute venous thrombosis. [1–3] There are all main drawbacks to this type of study. First of all, from the perspective of the venous thrombosis cycle, chronic thrombosis affects patients’ quality of life and treatment intervention, which can lead to venous ulcers and limb swelling.[4] Second, small rodents have a very different body size and hemodynamics than humans, which play a key role in flow-restricted, venous stasis-dominated thrombosis.[5] Finally, all the new technologies from basic research to clinical application, such as anticoagulant drugs, thrombolysis, thrombectomy devices, and endovascular treatment devices, cannot be studied with small animals.

Speaking of large animal models of venous thrombosis, there have been some attempts. The methods of these experiments are different, but the theories are inseparable from Virchow’s triad, including vessel wall injury, blood hypercoagulable state, and hemodynamic abnormalities. The surgical approaches used included open surgical procedures, endovascular techniques, and laparoscopic techniques.[6–8] Chantal Kang mentioned that 80% proximal stenosis and induced preoperative hypercoagulability could rapidly induce thrombosis and was highly consistent with histological morphology inside the human body. [9] This differs from thrombosis modeling in small animals, where small rodent thrombosis models can use simple proximal ligation of the inferior vena cava, which can rapidly form venous thrombus, whereas simple proximal ligation of the vein in large animals is difficult to form a thrombus, probably due to the huge blood flow and abundant collateral compensation in large animals.[10]

But so far, there is no long-term observation of a large animal model of chronic venous thrombosis, which requires at least 3 months and is consistent with chronic thrombotic obstruction of the iliofemoral vein in humans. To this end, we designed a model of the left iliac venous thrombus in Labradors and Bama miniature pigs. Because of the vascular anatomy, blood properties, and vascular physiology similar to humans, Labrador dogs and Bama miniature pigs were considered to be suitable experimental subjects. The present study reports the methods and follow-up results of this technique and the histological characteristics of the thrombus.

## Materials and Methods

### Animals

All adult Labrador dogs and Bama miniature pigs (provided by Taizhou Meifengli Animal Experimental Center, Jiangsu Province), weighed 20-30kg, male or female. All animal experiments passed the Guidelines for Ethical Review of Laboratory Welfare and followed the Regulations on the Administration of Laboratory Animals of the Ministry of Science and Technology, PRC. All animals were housed in single rooms at a temperature of 16-26°C and fed with standardized food and free access to food and water.

### Anesthesia

All animals were anesthetized with Zoletil (7-25 mg/kg,) intramuscular injection for sedation and maintained by endotracheal intubation during the operation with 1-6% isoflurane. Blood pressure, heart rate, electrocardiogram, and oxygen saturation were monitored during surgery.

### Induction of venous thrombosis in left iliac veins

In this model, we combined the stenosis model with thrombin to form an immediate and solid thrombus in the left iliac vein. Briefly, animals were placed in a supine position after anesthesia in a hybrid operating room. Conventional skin sterilization in abdominal surgery, a median abdominal incision was made from the first to the third nipple. Then we incised the skin and subcutaneous tissues until the rectus abdominis muscle and separated the lateral peritoneum bluntly alone with the left abdominal wall until the posterior peritoneum. Suspended the left iliac vein and artery with an absorbable thread after exposure of the abdominal aorta, inferior vena cava, and left common iliac artery and vein(Figure 1A). The circumference of the left common iliac vein was measured and the diameter of the common iliac vein was calculated(Figure 1B, C). The diameter of the ligating glass rod was calculated according to the degree of 80% stenosis(Figure 1D). The left common iliac vein near the bifurcation of the inferior vena cava was tied with a rod of the calculated diameter, and then the rod was pulled out to show the constriction of the iliac vein at the ligation site(Figure 1E). The diameter of the iliac vein after ligation was remeasured with silk thread again, and the stenosis rate of the model was calculated(Figure 1F, G). Then the first vascular clip was placed at the ligation site, and the second clip was used at the distal end 3 cm from the ligation site, taking care to avoid the branch veins(Figure 1H). The filling and dilatation of the common iliac vein were seen. After that, we injected sequentially 1 ml thrombin and fibrinogen with a 5 ml syringe under direct vision, and the blood flow was blocked with clips for 10 minutes(Figure 1I). Streptomycin 300 mg and ceftiofur sodium 300 mg were given postoperatively for 7 days. After 2 weeks of thrombosis model establishment, each animal was given oral treatment with rivaroxaban tablets at 20 mg per day.

**Figure 1:**
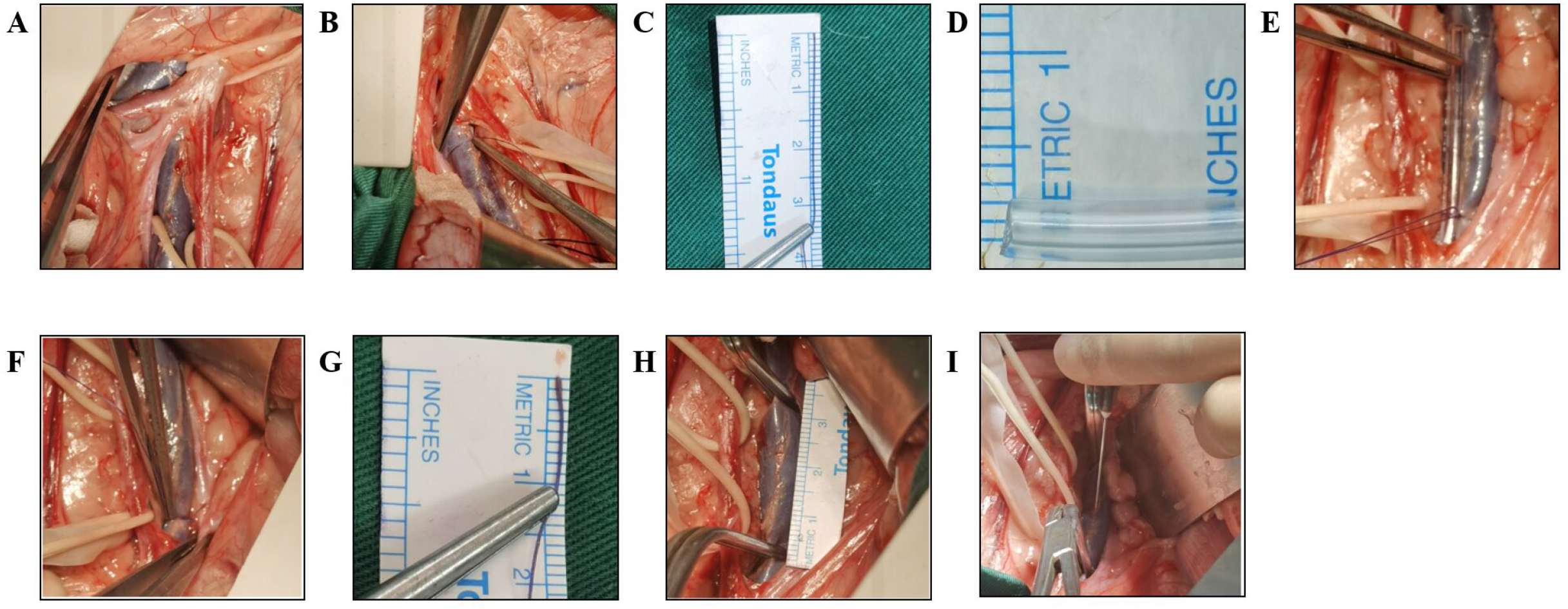
Induced Venous Thrombosis in Left Iliac Veins of Canines. A)Separated bilateral iliac veins. B, C)Measured iliac vein circumference. D)Diameter of the ligation rod. E) Ligation of rod and left iliac vein. F, G)Remeasured iliac vein circumference. El)Thrombosis length. I)Injection with thrombin and fibrinogen.

### Follow-up Venous Angiography

The left iliac vein blood flow was observed by anterograde angiography at 7, 30, 60, and 90 days after successful thrombus establishment. Under anesthesia described previously, the left inguinal area was disinfected and sheeted. The femoral vein was located by ultrasound and a 5F puncture sheath was placed into the femoral vein, and heparin water (8 U/mL, Shanghai ShangPharma First Biochemical Company) was injected through the catheter to prevent thrombosis from occluding the contrast catheter. Iodixanol contrast agent (100mL, General Electric Pharmaceutical Co.) and heparin water 10 ml (1:1) are pushed in for imaging. The left iliac vein should be visualized or the contrast agent should be returned to the inferior vena cava through the collateral vessels until the contrast agent dissipates in the visual field.

### Histological Staining

At 90 days after thrombosis, survived animals’ bilateral iliac veins were isolated and fixed in 4% paraformaldehyde for at least 24 h after dissection, followed by dehydration in ethanol with a gradient of concentration. The dehydrated samples were embedded in paraffin and serially sectioned at 4um thickness. These sections were subjected to HE and Masson staining and photographed by light microscopy.

## Results

All experimental animals completed left iliac vein thrombus formation successfully and there was extremely minimal surgical bleeding. Six Labrador dogs survived perioperative without intraoperative or postoperative pulmonary thromboembolism or death until 90 days while three Bama miniature pigs died on day 29, 33, and 58, respectively.

### The Canine Model of Chronic Deep Venous Thrombosis

The underlying condition of animals and operative data were shown in Table 1. The perimeter of dogs’ left iliac vein was 2.7-3.6 cm and the diameter was 0.86-1.15cm. Remeasurement of left iliac veins after ligation with a 0.2 cm diameter glass rod showed the constricted veins’ diameter was 0.29-0.48cm and the stenosis rates were calculated in Table 1. The length of each animal’s thrombus formation ranged from 3 to 3.5cm. In this process, we sequentially injected 1 ml of thrombin and fibrinogen under direct vision and occluded blood flow with vascular clamps for 10 minutes. Blood routine, blood biochemistry, and coagulation function were tested before the operation and 7, 30, 60, and 90 days after the operation. No evident abdominal wound bleeding or signs of hemogram infection occurred under postoperative anti-infection treatment.

**Table 1:** The underlying condition and intraoperative data of canines and pigs.

| Animal number | Canine number |  |  |  |  |  | Pig number |  |  | P value |
| --- | --- | --- | --- | --- | --- | --- | --- | --- | --- | --- |
|  | 1 | 2 | 3 | 4 | 5 | 6 | 1 | 2 | 3 |  |
| Sex | Male | Female | Female | Male | Male | Female | Male | Male | Female |  |
| Animal weight(kg) | 30.2 | 20 | 24.6 | 32.8 | 29.8 | 29.6 | 34.2 | 34 | 28.2 | 0.2046 |
| Duration of operation(min) | 78 | 62 | 51 | 88 | 87 | 80 | 147 | 109 | 175 | 0.0027(**) |
| Venous perimeter(mm) | 35 | 27 | 31 | 30 | 36 | 33 | 30 | 32 | 33 | 0.8773 |
| Venous diameter(mm) | 11.1 | 8.6 | 9.9 | 9.5 | 11.5 | 10.5 | 9.5 | 10.2 | 10.5 | 0.8664 |
| Remeasured venous perimeter(mm) | 11 | 9 | 14 | 15 | 12 | 15 | 15 | 12.0 | 14 | 0.5415 |
| Remeasured venous diameter(mm) | 3.5 | 2.9 | 4.5 | 4.8 | 3.8 | 4.8 | 4.8 | 3.8 | 4.5 | 0.5489 |
| Stenosis rate | 90.12% | 88.89% | 79.60% | 75.00% | 88.89% | 79.34% | 75.00% | 85.94% | 82.00% | 0.5623 |

Clinically, patients with chronic iliofemoral vein thrombosis have angiographic manifestations of vein occlusion or smaller caliber, with significant collateral circulation and distal dilation. All animals were accepted for antegrade venography of left iliac veins at follow-up time points to observe the formation of thrombus and the restriction of blood flow. As shown in Figure 2, 7 days after modeling, no contrast material passed through the left common iliac vein in all animals. A small amount of collateral circulation was observed, the thrombus segment was visible and the distal vessel was distended. At 30 days, angiography of the thrombus showed discontinuous filling defects or narrowing of the lumen. The possible cause was thrombolysis and the thrombus organization because of postoperative anticoagulation drugs, which attached to the vein wall and caused blood flow restriction. Collateral compensation was significantly increased at 60 days compared with 30 days after modeling, and distal limb blood flow returned to the heart from the collaterals. At 90 days, the collateral circulation gradually stabilized, the collateral vessels became thicker, and the blood flow was dominated by several major collateral branches.

**Figure 2:**
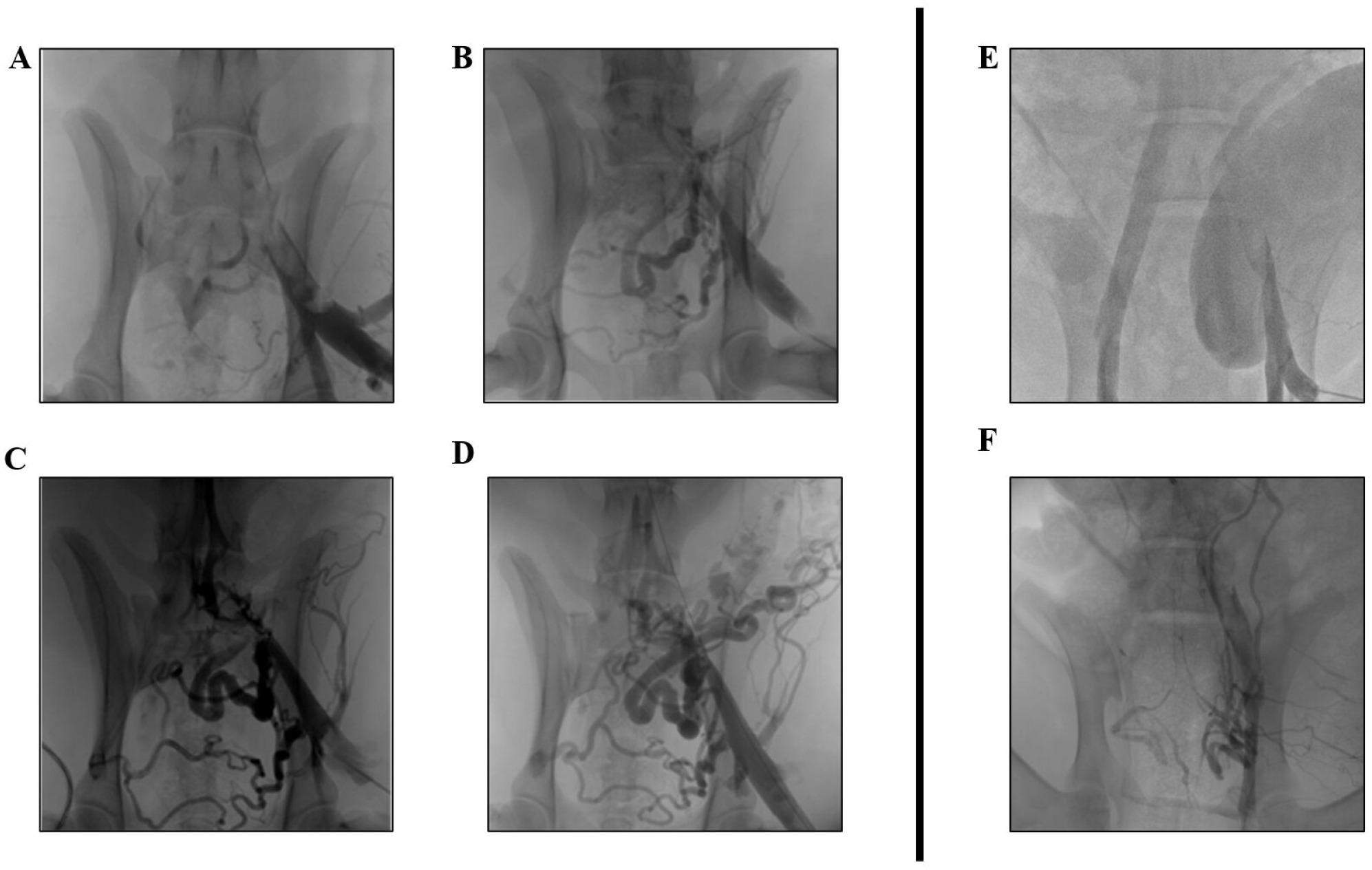
Venous Angiography. A-D: 90 Days Follow-up Venous Angiography of Canines. A) Days 7. B) Days 30. C) Days 60. D) Days 90. E_x_ F: 30 Days Follow-up Venous Angiography of Pigs. E) Days 7. F) Days 30.

Histological staining of bilateral iliac veins was performed on day 90 of thrombosis, and a large number of erythrocytes, inflammatory cells and fibroblasts were seen in the thrombosed segment(Figure 3). The platelet structure composed of collagen fibers was surrounded by layers with clear boundaries and interspersed with a large number of scattered erythrocytes, and a large number of lymphocyte infiltration and fibroblasts were seen in each platelet layer. In addition, the endothelial layer of the vein was damaged, multiple endothelial cells were seen to be lost, and the vessel wall was thickened.

**Figure 3:**
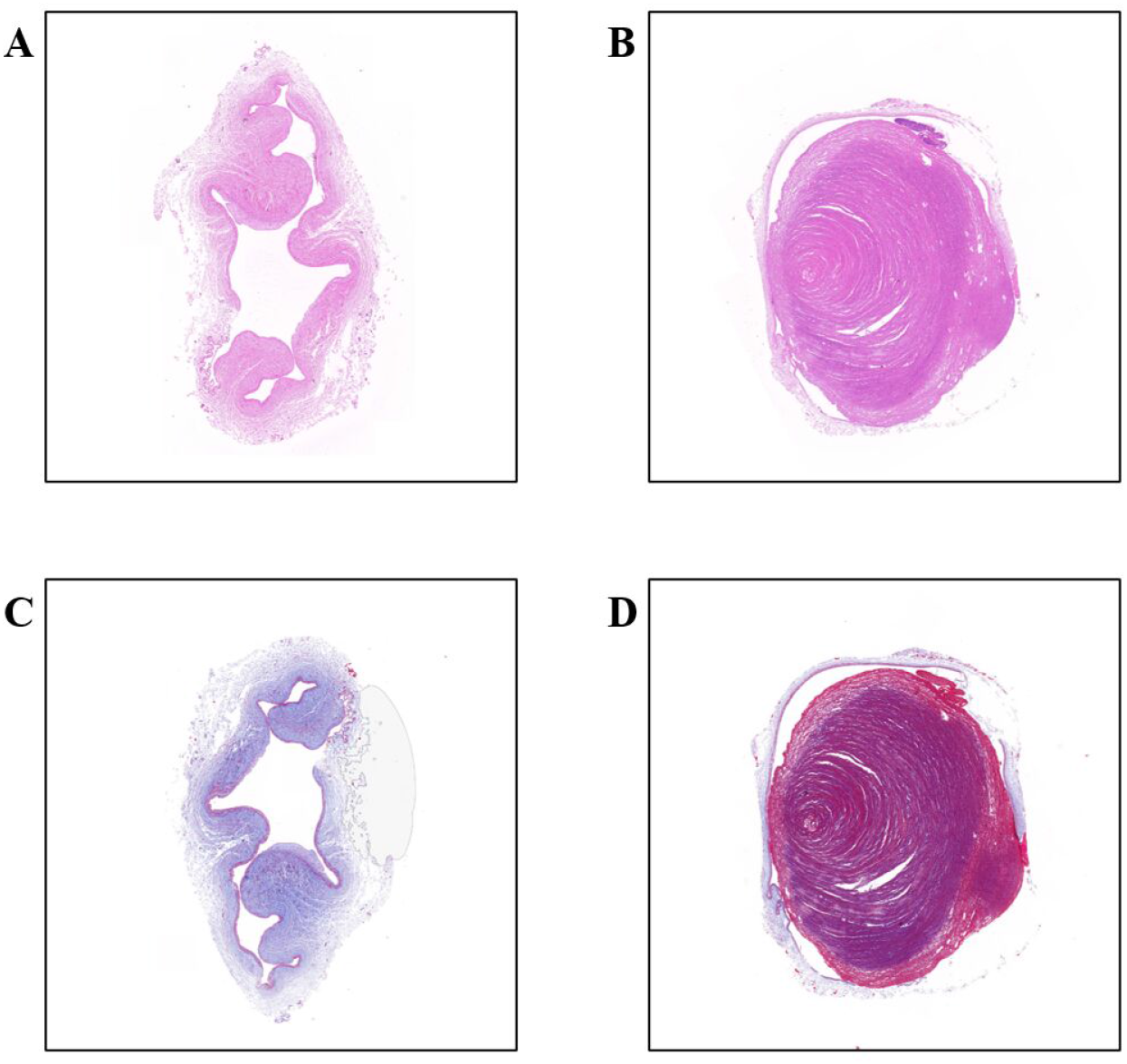
HE and Masson stain of bilateral iliac veins 90 days after modeling in dogs. A)HE stain of right iliac veins. B) HE stain of left iliac venous thrombosis. C) Masson stain of right iliac veins. D)Masson stain of left iliac venous thrombosis.

### The Porcine Model of Chronic Deep Venous Thrombosis

For some reason, three Bama miniature pigs died on day 29, 33, and 58 after venous thrombosis formation, respectively. The animal monitors did not see any noticeable abnormalities during daily inspections until the animals were found to die suddenly, without any signs before death, and the surgical sites were incised with tight skin sutures and healed well. In Table 1, there was no significant difference between canines and pigs except for the duration of surgery. Time of thrombus modeling in pigs was much longer than dogs for pigs’ complex lower abdominal and pelvic anatomy(Figure 4). The inferior vena cava and bilateral iliac veins of pigs were in the fat-filled retroperitoneal tissues, affecting vascular separation and exposure(Figure 4A). The diameter and length of left iliac veins were similar and the surgical procedure was consistent with dogs. Furthermore, postoperative follow-up angiography revealed deficient collateral compensatory capacity. In Figure 2F, at 30 days, only a few and tiny lateral branches appeared. Autopsy reports of three animals suggested thoracic infection, tissue bruising and pulmonary hemorrhage as the cause of death(Figure S1).

**Figure 4:**
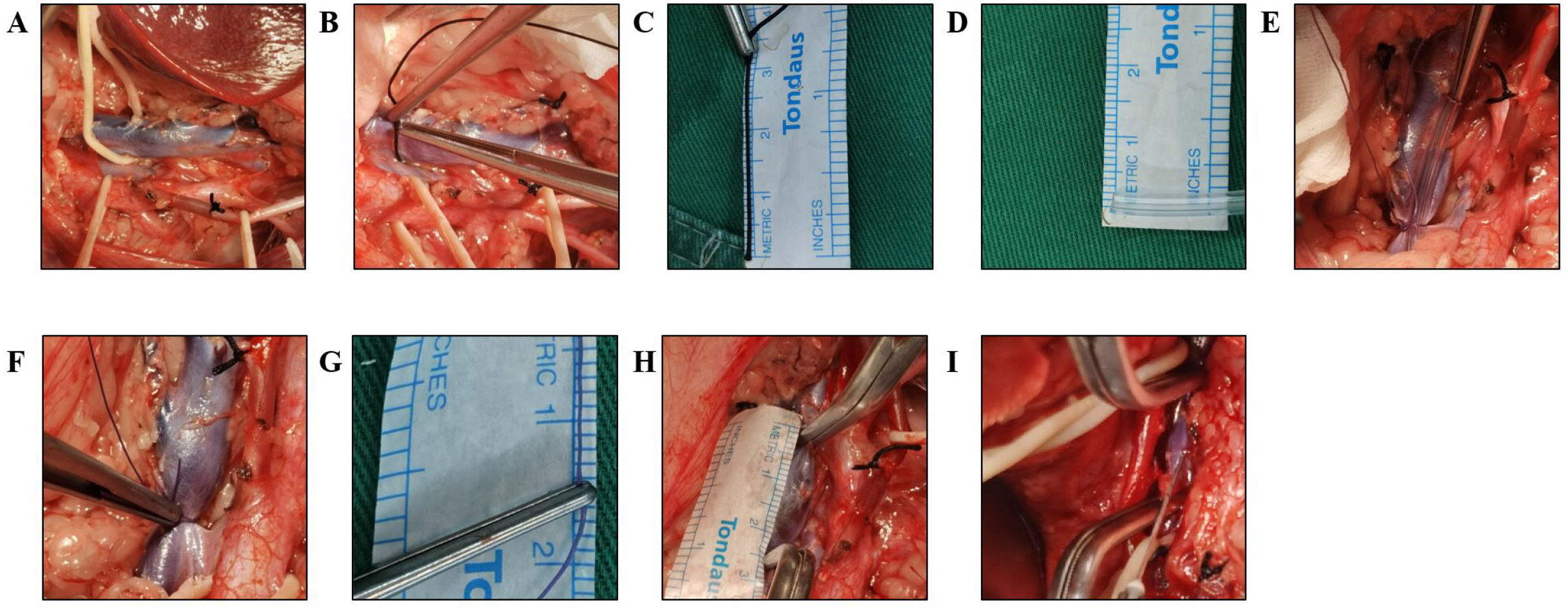
Induced Venous Thrombosis in Left Iliac Veins of Pigs. A)Separated bilateral iliac veins. B, C)Measured iliac vein circumference. D)Diameter of the ligation rod. E) Ligation of rod and left iliac vein. F, G)Remeasured iliac vein circumference. H)Thrombosis length. I)Injection with thrombin and fibrinogen.

## Discussion

Deep venous thrombosis is a common disease in clinics. [11] Establishing animal models that conform to the characteristics of human thrombosis is a necessary method to reveal the pathogenic mechanism, research drugs, and devices improve treatment strategies. Small animal models, especially rodents, are the most commonly used research tools. However, due to their size differences, small animal models cannot be used for translational research of various medical devices, which limits the development of new interventional or surgical open treatments. Almost half of the large animal thrombosis studies have chosen pig as the large animal model for venous thrombosis research for pig has similar vascular anatomy and size to human, to test therapeutic drugs and devices, to test surgical procedures, to assess DVT imaging, to study DVT pathophysiology.[12] However, a load of pelvic and lower limb blood reflux due to an inferior cavity or iliofemoral vein thrombosis makes pressure-sensitive pigs susceptible to death, with a mortality rate of up to 33%.[13] Compared with other large experimental animals such as pigs or sheep, dogs are more compliant and easier to handle during anesthesia and follow-up, while they also have stronger resistance to infection and higher survival rates. More importantly, in the chronic phase of human DVT, it is emphasized to increase the amount of activity to avoid the spread and recurrence of thrombosis caused by blood stasis.[14] The active and active nature of dogs is more consistent with the development law of chronic thrombosis in the human body. The diameters of veins used in DVT models varied from 6.3 mm to 14 mm reported rarely.[12] In this study, the diameter of the left common iliac vein in Labrador dogs was 8.6-11mm, which was close to the diameter of the human iliac vein, and closer to the human body when studying interventional medical devices.

The most common method of modeling venous thrombosis is the venous stasis model, in which blood flow is restricted or temporarily blocked by ligation of varying degrees of stenosis, balloon blockade, or cone stenting[15], and local thrombus formation is induced by exogenous injection of thrombin. There is also significant thrombus formation in the lumen by stimulation of the vascular endothelium by anodal direct current.[16] Notably, in models using the intraluminal technique, blocking flow using both ends of the balloon and thrombin catheter administration to promote thrombus formation, the balloon block time was maintained for at least 1 hour, regardless of vessel diameter (jugular, iliac, inferior vena cava).[17] The time required was longer using the electrical stimulation method.[16] In this experiment, the innovative use of both thrombin and fibrinase resulted in solid thrombus formation visible within about 5-7 minutes of the intraoperative blockade, greatly reducing the time of the modeling procedure and mitigating the risk of anesthesia in the animals.

The canine model of chronic thrombosis in this study recapitulates the features of post-thrombotic syndrome in humans. Left iliac vein outflow obstruction due to hemodynamic abnormalities and thrombosis is the main cause of recurrent episodes and poor prognosis in patients with clinical chronic thrombosis. In addition, this study is the first to use angiography to observe thrombosis for up to 90 days, providing a model consistent with chronic thrombosis progression. Furthermore, the measurable stenosis rate of venous obstruction can be divided into different groups to determine the timing of intervention.

This model can be used to study the biological safety and efficacy of anticoagulant drugs, thrombectomy, thrombolytic devices, and various new venous stents for venous obstruction. We compared the ability of pigs and dogs to tolerate long-term chronic thrombosis and showed that pigs died suddenly due to thoracic infection, tissue bruising, and pulmonary hemorrhage. Possible triggers were prolonged blood flow obstruction due to massive thrombosis of the common iliac vein, including restricted internal iliac vein return, poor collateral compensation, and severe pelvic stasis, which induced gastrointestinal stress and leaded to malnutrition and infection. In addition, the pigs may have poor sensitivity to anticoagulant drugs and susceptibility to bleeding risk, which also influenced the success of the model. More than that, dogs have superior health, obedience, and activity than pigs and this may be the reason why the canine model is more successful.

## Highlights

1. This study is the first innovative model of chronic thrombosis in dogs and pigs, with an observation period of up to 90 days and a mature and reliable modeling method.
2. Canines have a higher safety profile for use in chronic deep venous thrombosis models due to their clear anatomy, excellent obedience, and animated behavior compared with pigs.
3. In order to progress in endovascular luminal techniques and device research, it is indispensable to adopt large animal thrombosis models that are more in line with the performance within the human body.

## Contributions

Chuang Wang, Tao Tang and Sheng-Lin Ye designed and implemented this animal model, Xu-Dong Jiang, Guang-Yuan Xiang and Lu-Lu Wang were responsible for data collection, and Tian-Ze Xu, Bin Song and Han Kang analyzed data. Chuang Wang wrote this paper.

## Acknowledgments

This work was supported by the National Natural Science Foundation of China (82070496, 81770483, 82100517, 32101104), the Natural Science Foundation of Jiangsu Province (SBK2020040321), the China Postdoctoral Science Foundation (2020M670035ZX), the Fundamental Research Funds for the Central Universities (0214-14380481), the Postdoctoral Research Funding Program of Jiangsu Province (2020Z368), the Nanjing special fund for health science and technology development (YKK21073), and the 2020 Jiangsu Province Shuangchuang Ph.D. Introducing Talent Project of Nan Hu.

---

**Supporting Figure 1:**
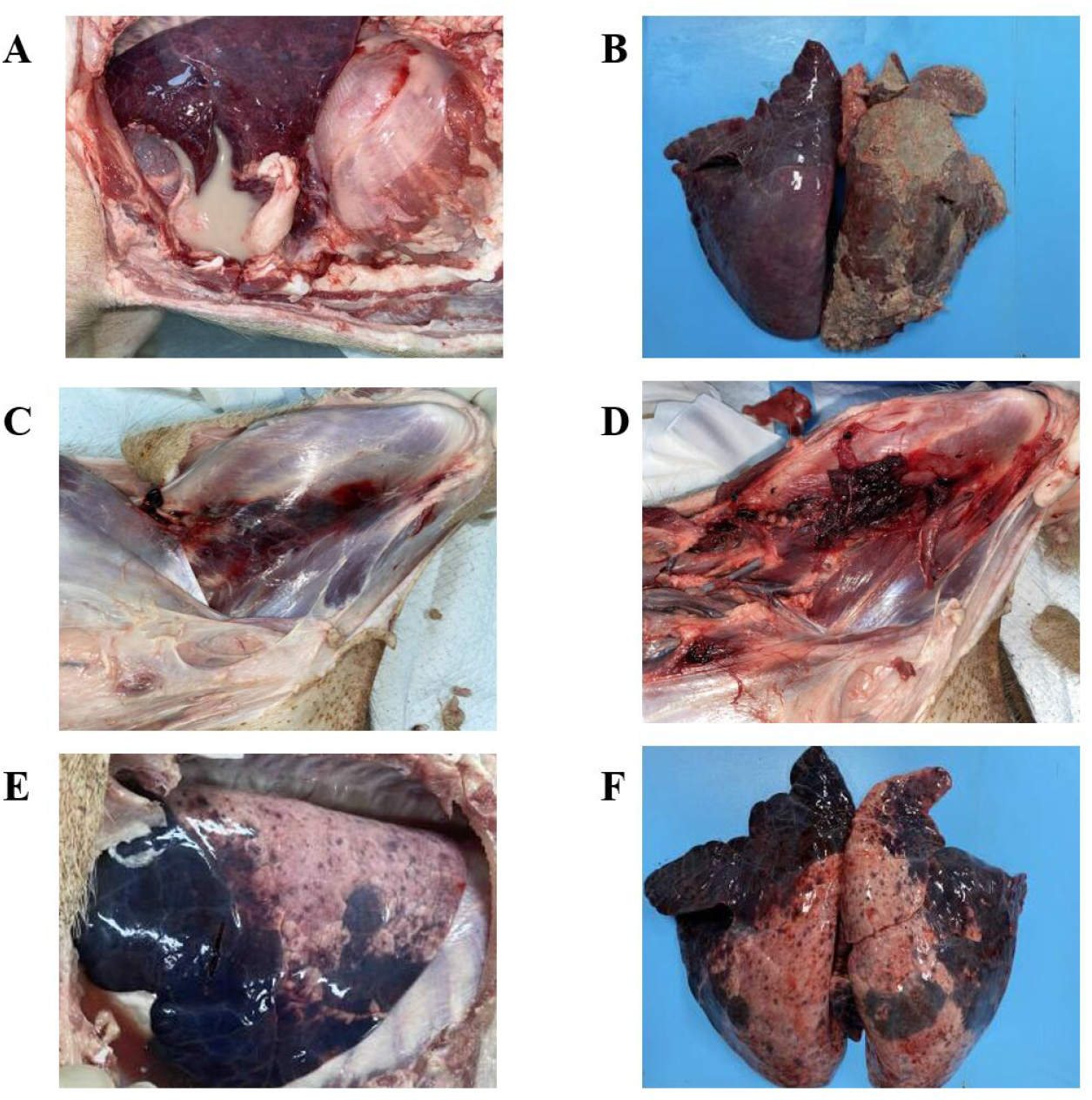
Autopsy reports of three Bama miniature pigs. A, B: 29 Days. C, D: 33Days. E, F: 58 Days

